# Tomographic Printing in a Chip: A Versatile Platform for Biomimetic 3D Organ-on-Chip

**DOI:** 10.64898/2026.02.26.708161

**Authors:** Riccardo Rizzo, Viola Sgarminato, Felix Wechsler, Christophe Moser

**Author notes:** Funding: Swiss National Science Foundation Return CH Postdoc.Mobility fellowship P5R5-3_235066 (RR) Swiss National Science Foundation under grant number 10007068 “Neural precision Holographic Volumetric Additive manufacturing” (FW).

## Abstract

Organ-on-chip (OoC) platforms are increasingly adopted for predictive in vitro testing. However, most remain limited by soft-lithography–derived 2.5D microfluidic architectures and non-physiological rigid materials, or bioprinting approaches that require complex and failure-prone post-fabrication assembly. Here, we present a versatile approach that integrates tomographic volumetric additive manufacturing (TVAM) directly within preassembled microfluidic chips, enabling rapid, contactless fabrication of freeform 3D OoCs. Leveraging our open-source optical simulation framework, Dr.TVAM, we perform TVAM in custom-designed chips, eliminating post-printing manual assembly steps that commonly lead to leakage, contamination, and poor reproducibility. This strategy, termed TVAM-in-a-chip, supports the generation of diverse 3D channel architectures in multiple biocompatible photoresins spanning a wide range of chemistries and mechanical properties, including cell-laden formulations. We demonstrate multi-channel designs, compatibility with confocal imaging, and dynamic culture of epithelial and endothelial models. Overall, TVAM-in-a-chip overcomes key limitations of current OoC technologies and paves the way for a new generation of scalable, biomimetic 3D platforms for advanced in vitro modeling.

## 1. Introduction

Human organ-on-chip (OoC) platforms are an emerging technology with broad applications in drug discovery, chemical and food testing, and to study environmental impacts on human health.^1,2^ OoCs are defined as “microphysiological systems that replicate one or more aspects of an organ’s in vivo dynamics, functionality, and/or (patho)physiological responses”.^3^ Over the past decade this technology has experienced a dramatic growth which led to the development of numerous tissue models.^2^ Pharmaceutical companies and regulatory agencies, including the FDA and EPA, have begun to adopt organ-on-chip (OoC) platforms for safety and efficacy testing,^4^ thereby complementing and reducing reliance on animal testing.

To date, OoCs mostly consist of microfluidic devices manufactured via soft lithography.^5,6^ On one hand, this well-established approach is advantageous as it is relatively inexpensive and offers high-throughput fabrication. However, it results in overly simplified devices composed of non-biomimetic rigid materials (e.g., PDMS) with 2.5D perfusable geometries confined to a single plane and featuring non-physiological architectures and square cross sections **(Figure 1A)**. These aspects limit OoC biomimicry, and thus their potential clinical predictivity.

**Figure 1.**
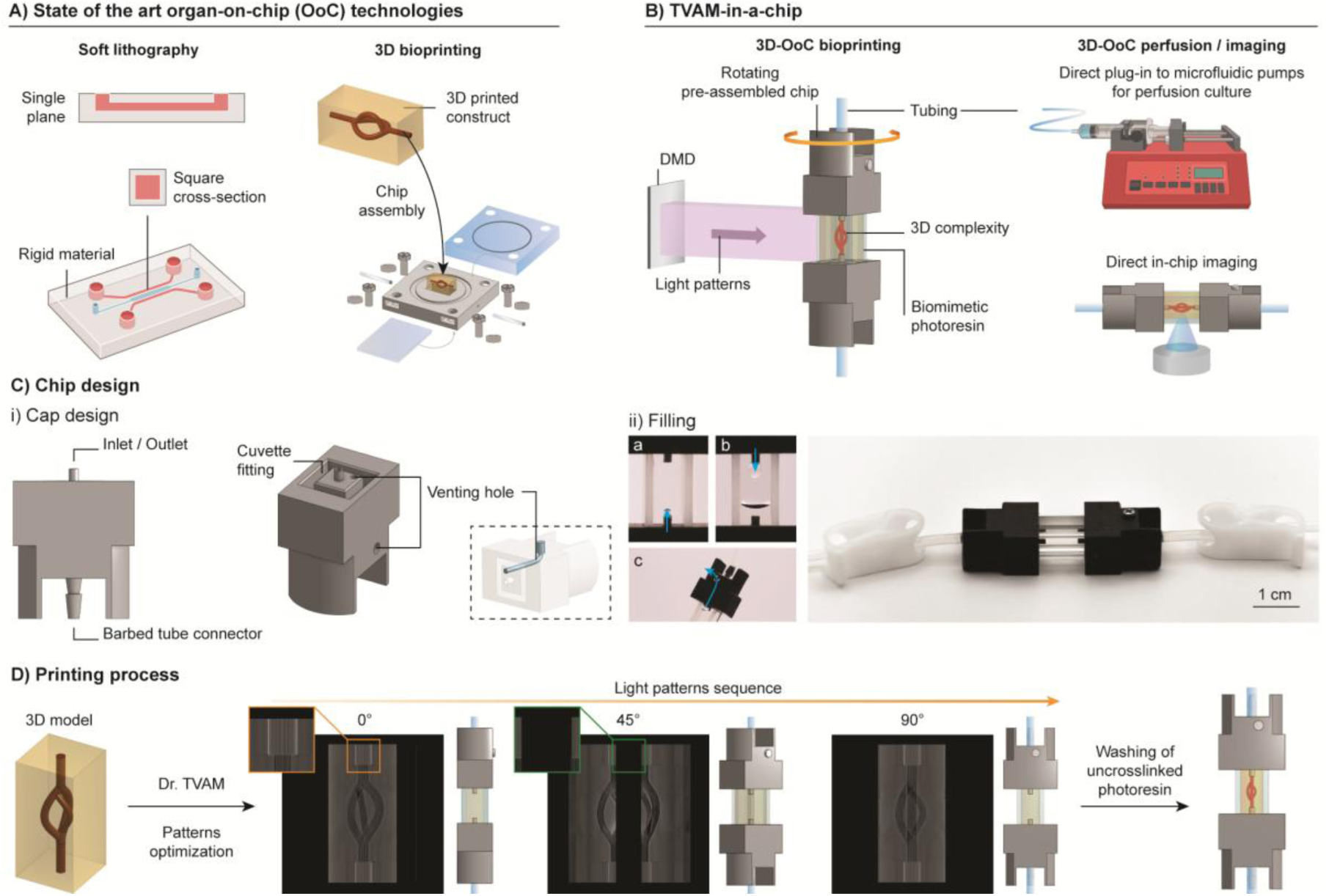
Rationale, and working principle of TVAM-in-a-chip for OoC applications. (A) Schematic of soft-lithography-based and bioprinting-based OoCs. (B) Schematic of TVAM-in-a-chip. (C) Details of custom chips (i), filling steps and fully assembled chip (ii). (D) Schematic of printing process with examples of intensity projection patterns at three rotation angles. Orange and green close-ups highlight regions of higher intensity at the light-occluding inlet (due to overprinting condition) and zero intensity at the cuvette corner to prevent scattering, respectively.

In the effort to implement biological matrices and organotypic microarchitecture to OoCs, in recent years several groups have explored various 3D bioprinting approaches such as embedded printing,^7–9^ and digital light processing.^10,11^ Overall, these methods offer ways for enhanced organ mimicry across multiple length scales. However, several challenges remain. In addition to the lengthy printing processes which limit their scale up and throughput, a key issue of current technologies is represented by the multiple post-printing steps needed to assemble the chips which lead to a significantly increased risk of failure **(Figure 1A)**. For example, embedded printing is conducted in an open chip, which is ultimately manually sealed. For DLP, traditional extrusion printing and other approaches, the printed models have to be post-processed, often manually handled, transferred to an open chip which is only then sealed and connected to a microfluidic system. For those engaged in the development of such OoCs, issues such as leakage, contamination, and poor reproducibility are not uncommon.

In recent years a novel light-based printing method named tomographic volumetric additive manufacturing (TVAM) has gained a lot of interest for its unique ability to generate complex centimeter-size models in seconds.^12–14^ In this layer-less printing technology, pre-calculated 2D projection patterns are sent towards a rotating vial filled with a photosensitive resin (photoresin). This process results in a 3D light dose accumulation in the photoresin. When the dose exceeds the gelation threshold, the desired model is formed. A few studies have begun to explore the use of TVAM to generate perfusable structures and tissue models.^15–17^

Although extremely fast and versatile in terms of attainable model complexity, this approach still suffers from the need to remove the printed construct from the round vial for post-processing and assembly in a chip with flat surfaces for imaging purposes. Recently, our group developed an open-source optical framework (Dr.TVAM) which could tackle this unmet challenge.^18,19^ This differentiable, physically-based ray-optics approach supports TVAM in arbitrary geometries, such as square vials, and in overprinting scenarios (presence of light occluding elements). In this work, we leverage this simulation software to perform contactless TVAM printing of freeform 3D organotypic perfusable models within preassembled, leakage proof chips with integrated fluidics **(Figure 1B)**. This approach, named TVAM-in-a-chip, retains the great versatility of TVAM, thus enabling the generation of chips with diverse 3D perfusable geometries using a variety of biocompatible photoresins that differ in chemistry (polymer backbone, crosslinking mechanism, and photoinitiating system) and mechanical properties (viscosity and stiffness). We further demonstrate TVAM-in-a-chip versatility by printing chips with up to three input-output channels, and with cell-laden resins. Importantly, given their flat surface and miniaturized size, such OoCs are compatible with standard imaging methods, allowing for direct in-chip confocal microscopy of stained samples throughout the whole volume. Finally, as proof-of-concept, we further demonstrate successful seeding and culture of epithelial and endothelial cells in such devices.

The proposed strategy emerges as a versatile tool to overcome the limitations of current 2.5D chips and build off the potential of TVAM, paving the way for more complex biomimetic 3D OoC platforms.

## 2. Results

### 2.1. TVAM-in-a-chip

First, we designed miniaturized custom chips compatible with TVAM. The optical clear container with flat surfaces was obtained from 5 mm wide square quartz cuvettes cut into 20 mm long sections. This cuvette section, which represents the main chamber of the chip, is then sealed on both sides with SLA biocompatible printed caps. The caps were designed to feature a 5 mm hole to press-fit the cuvette, leaving a printable window of 10 mm. Caps also featured a 1.5 mm inlet with 0.8 mm internal channel, a round holder for mounting on the rotating stage, and a barbed connector to interface the device with microfluidic tubing (Fig. 1C-i, and Fig. S1). The chip, thus composed of the main quartz chamber, two caps, and tubing is then filled with photoresin. To eliminate large bubbles that could affect the printing process, both sides are primed via photoresin injection through the tubing. On one side only the cap also includes a venting hole to ensure that, upon filling the chip with photoresin, bubbles could be evacuated **(Figure 1C-ii)**. Finally, the venting hole is sealed with a screw, and the preassembled chip is ready to be used for the TVAM process **(Figure 1C-ii)**. Importantly, the footprint of the proposed miniaturized chips is on par with that of commercially marketed soft-lithography-based platforms.

The projection patterns were calculated using Dr.TVAM,^18^ and accounting for a number of parameters including shape and refractive index of the cuvette, absorption and refractive index of the photoresin, but also shape, position and optical properties of the occluding elements (inlets). The inlets, fabricated with a black resin, were assumed to be fully absorbing. To counterbalance this light absorption along the light path, Dr.TVAM puts more intensity in these regions, ensuring a homogeneous print (**Figure 1D**, orange close up). The corners of the cuvette were discarded from the optimization (zero intensity regions) to avoid unwanted scattering (**Figure 1D**, green close up). Once mounted to the printer stage, the projections patterns (600 per turn) are synchronized with the rotation of the chip, as for a standard TVAM process. When the desired model is printed, the uncrosslinked resin is washed out by perfusing the microfluidic system. Importantly, as the printing happens within the chip, no postprocessing step is required to take out the construct. The OoC is then ready to be used for further experiments, such as cell seeding and dynamic culture via connection to microfluidic pumps, as later shown.

### 2.2. Versatility across multiple photoresin formulation

A key advantage of TVAM-based systems lies in their potential compatibility with diverse biocompatible matrices. To test this hypothesis, we investigated the compatibility of TVAM-in-a-chip with synthetic (4-arm PEG methacryloyl, PEG4-MA), polysaccharide (hyaluronan-methacryloyl, HA-MA) and polypeptide (gelatin-methacryloyl, Gel-MA) derived resins. These resins, all incorporating 0.05% LAP as photoinitiator, were used at different concentrations to attain similar reactivity (**Figure 2A**). Notably, the 20% PEG4-MA, 2% HA-MA and 10% Gel-MA result in significantly different storage modulus (G’) under photorheology measurement, respectively (85.51 ± 2.19) kPa, (4.73 ± 0.04) kPa, and (1.99 ± 0.08) kPa (**Figure 2A**). As shown in **Figure 2A**, all three resins could be successfully used to generate perfusable spiral structures in a chip. In these and future examples, the printed shape is perfused with red-, green-or blue-dyed water-based solutions for better visualization. Interestingly, despite showing similar gelation onset in photorheology, the resins required significantly different light doses, going from 31.1 mJ cm^-3^ for PEG4-MA to 11.6 mJ cm^-3^ for HA-MA and 6.3 mJ cm^-3^ for Gel-MA. This difference can be linked to the resin viscosity. In standard TVAM processes, an optimal resin viscosity greater than 10 Pa·s is often reported to prevent sedimentation.^12^ In contrast, the PEG4-MA and HA-MA formulations used here exhibit substantially lower viscosities: (5.1 ± 2.0) mPa·s for 20% PEG4-MA and (120 ± 17) mPa·s for 2% HA-MA. 10% Gel-MA is instead allowed to thermally gel at room temperature prior to printing. For PEG4-MA and HA-MA resins, due to the low viscosity, diffusion of oxygen (inhibitor) and radical species still play an important role in delaying crosslinking. However, because the cuvette is sealed and printing is performed wall-to-wall, issues such as sedimentation of ongoing prints does not occur, enabling the use of very low-viscosity resins in TVAM-in-a-chip.

**Figure 2.**
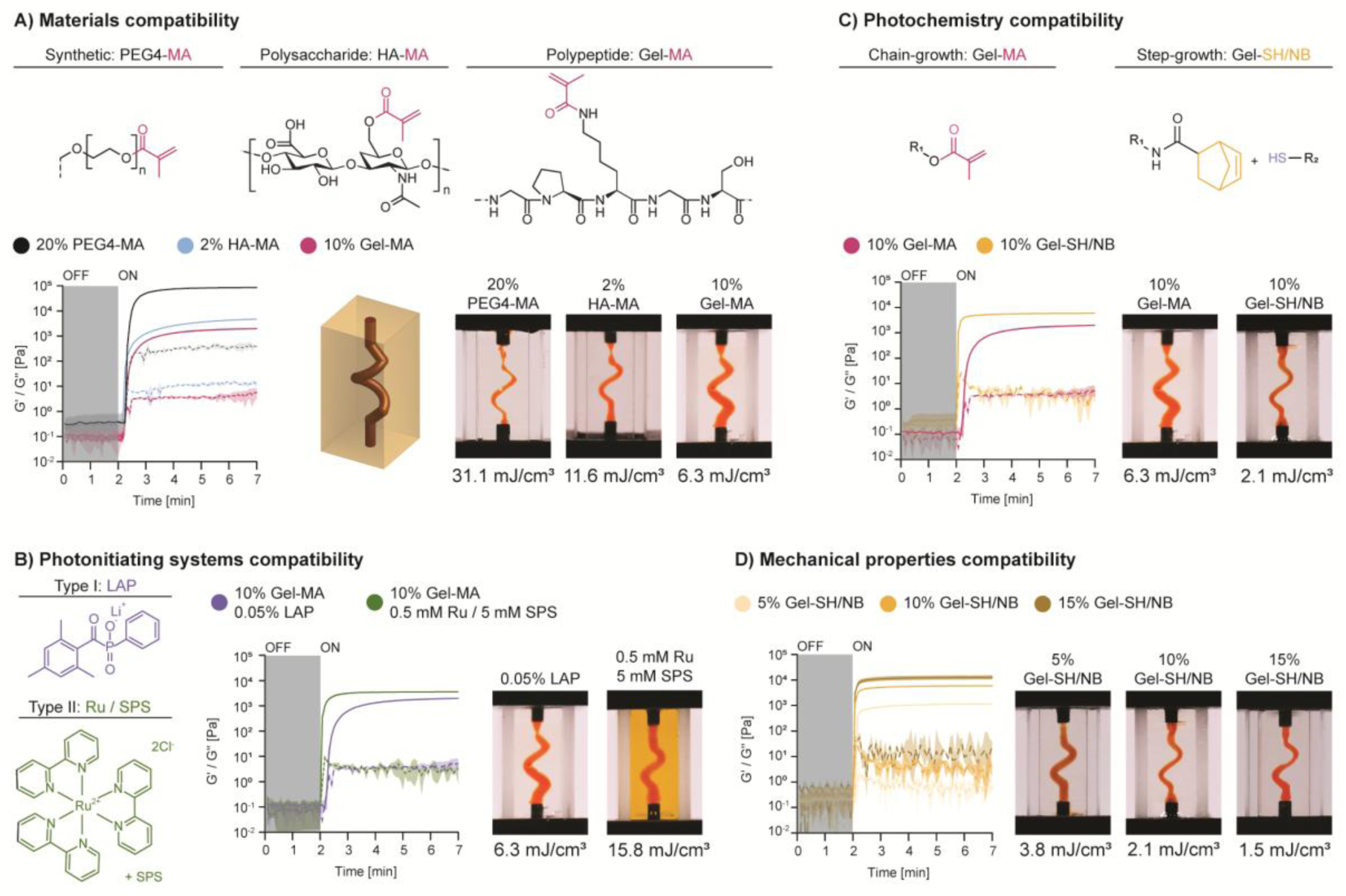
TVAM-in-a-chip compatibility with diverse photoresin chemical and mechanical properties. (A) Chemical structure, photorheology and spiral prints of synthetic, polysaccharide and polypeptide-derived photoresin formulations. (B) Chemical structure, photorheology and spiral prints of gelatin-based chain-growth and step-growth photoresins. (C) Chemical structure, photorheology and spiral prints of gelatin-based step-growth photoresins with Type I and Type II photoinitiators. (D) Photorheology and spiral prints of gelatin-based step-growth photoresins at various polymer concentrations.

Although LAP, a Norrish type I initiator, became the gold standard for bioprinting purposes for its high-water solubility and reactivity, other systems have been explored. Among these, a Norrish type II initiating system composed of Tris(2,2′-bipyridyl)dichlororuthenium(II) and sodium persulfate as co-initiator (Ru / SPS) has gained particular attention for its biocompatibility and applicability to (meth)acrylate-based resin,^20,21^ and tyrosine-based resins.^22^ In this work, we show that TVAM-in-a-chip is compatible with both Type I and Type II initiating systems (**Figure 2B**), thus further expanding the palette of chemistries and materials that can be adopted. Interestingly, although photorheology measurements indicate faster curing kinetics for the Ru / SPS formulation at the tested concentrations, printing required more than twice the exposure dose compared with the LAP formulation (15.8 vs 6.3 mJ cm^-3^). This behavior reflects the limited stability of SPS in water-based, buffered polymer solutions. During the few minutes required for thermal gelation in the chip, SPS is chemically consumed by oxidizing both amino acid residues in gelatin and the ruthenium complex Ru(II) to Ru(III). Accordingly, it is generally advised to prepare SPS freshly, add it last to the formulation, and minimize the time the resin remains in the dark prior to irradiation.

We then investigated the compatibility of our approach with different crosslinking chemistries, namely chain-growth (methacryloyl-based) and step-growth (thiol-norbornene based). Thiol-norbornene chemistry is known to be among the fastest biocompatible crosslinking strategies, also resulting in improved network homogeneity and mechanical properties when compared to methacryloyl systems.^15,23,24^ As expected, both photorheology and printing show a significantly faster crosslinking for the 10% gelatin-thiol / gelatin-norbornene (Gel-SH/NB) formulation compared to 10% Gel-MA. In particular, 10% Gel-SH/NB requires 3-times less light dose than 10% Gel-MA (2.1 vs 6.3 mJ cm^-3^).

Finally, we show that one can tailor the resin concentration to target different stiffness values (**Figure 2D**). In this example we successfully printed with 5%, 10% and 15% Gel-SH/NB which resulted in storage moduli of (1.15 ± 0.05) kPa, (5.90 ± 0.25) kPa, and (12.83 ± 1.85) kPa, respectively. As expected, the required light dose to cross the gelation threshold decreases with higher polymer concentration: 3.8, 2.1 and 1.5 mJ cm^-3^ for 5%, 10% and 15% Gel-SH/NB, respectively.

Altogether, these prints demonstrate the compatibility of TVAM-in-a-chip with a broad range of biocompatible resins spanning diverse (photo)chemical and mechanical properties, thereby defining a wide design space for the development of tissue-tailored OoC matrices.

### 2.3. Design flexibility and imaging capabilities

In addition to resin versatility, TVAM-in-a-chip also offers significant flexibility in design. The caps can be further customized to incorporate up to two or three inlets. Consequently, we demonstrate the ability to print perfusion systems composed of one, two, or three channels connecting inlets on opposite sides (**Figure 3A-i**). Moreover, caps with different inlet numbers can be combined within the same chip, thus also providing the possibility to print channels with inlet and outlet on the same side or mixing components converging the flow from two or three inlets into a single outlet (**Figure 3A-ii**).

**Figure 3.**
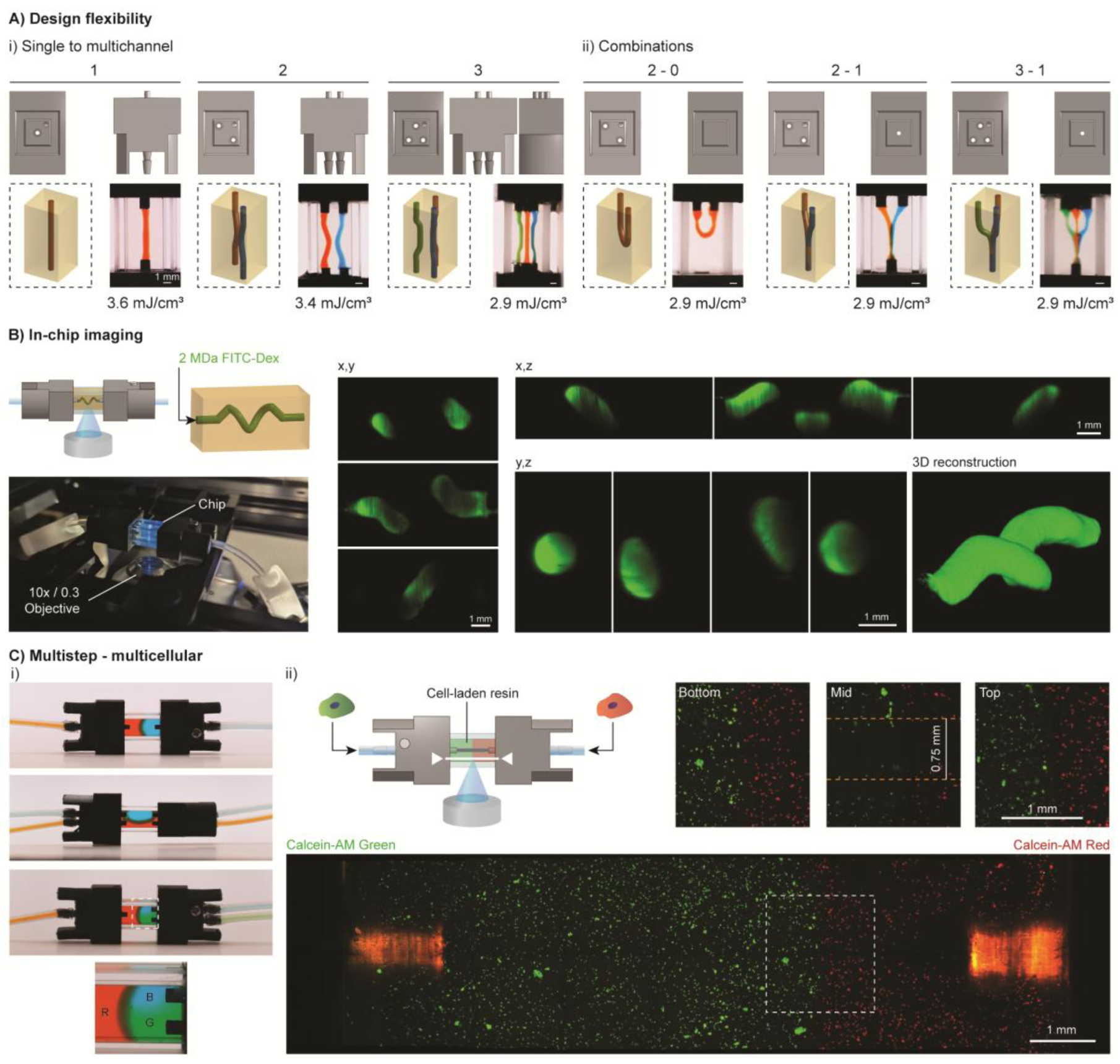
Design flexibility and imaging capabilities in TVAM-in-a-chip. (A) CAD views of caps (top), 3D model (dashed box) and pictures of perfused (red, blue, green) printed channels in chips featuring the same inlet/outlet numbers (i) or their combinations (ii). (B) Schematic of a chip featuring a spiral channel perfused with 2 MDa FITC-Dextran (top-left). Picture of the chip during confocal imaging (bottom-left). Confocal images of fluorescent spiral from different views and 3D reconstruction. (C) (i). Examples of step-wise filling of various chips with colored resin (5% Gel-SH/NB). (ii) Schematic of multistep filling with cell-laden resin featuring green- and red-labeled cells (top left). Confocal image of the cell-laden chip (white line in the schematic indicates the reference position). High-magnification views (top right) highlighting the interface region showing different cell populations and channel at varying depths. Note: red autofluorescence of the two inlets on both sides.

We initially chose to adopt a miniaturized cuvette with 5 mm path length not only to minimize the footprint of the chip, but also to guarantee access for direct (in-chip) confocal imaging throughout its volume. To verify this hypothesis, we printed a spiral shape in 5% Gel-SH/NB resin and perfused it with high-molecular weight FITC-Dextran (2 MDa). The custom chip fits into standard confocal stages (**Figure 3B**) and can be imaged with a long working distance objective (10x Mag, 0.3 NA, 11 mm WD). The full chip volume was accessible for scanning, allowing imaging of the complete 3D perfusable geometry.

Finally, we further explored the system versatility by performing stepwise filling of colored resin in different chip arrangements (**Figure 3C-i**), showing the possibility to introduce multiple properties in different locations. As a practical example, we filled a chip with green-labeled human foreskin fibroblasts (HFFs) and red-labeled HFFs on two different sides (**Figure 3C-ii**). We then performed the TVAM process in the cell-laden chip, generating a straight channel connecting the two inlets. Upon perfusion with PBS to wash out the uncrosslinked resin and embedded cells in the channel, we imaged the chip via confocal microscopy. As shown in the different imaging planes in **Figure 3C-ii**, live green- and red-labeled cells are located in distinct regions of the chip, while no cells are observed in the central area where the 0.75 mm channel was printed. Notably, this example shows the possibility to perform TVAM-in-a-chip in the presence of cells and further demonstrates our ability to achieve full imaging depth within the chip.

### 2.4. Reproducibility, resolution and complexity

To evaluate the reproducibility and scalability of the rapid TVAM-in-a-chip approach, five chips incorporating a spiral channel were fabricated in five consecutive prints using an identical light dose (3.8 mJ cm⁻³) **(Figure 4A)**. Given the printing parameters used for this model (see **Table S1**) and the roughly 30 seconds to mount and align each cheap, the printing process was completed in about 5 minutes. Channel diameters were measured at multiple positions to assess both accuracy and precision. The central regions of the structure showed high fidelity to the design, with the first turn (0.72 ± 0.02 mm), middle section (0.70 ± 0.03 mm), and second turn (0.69 ± 0.03 mm) closely matching the target diameter of 0.7 mm and exhibiting minimal variability across prints. In contrast, regions near the inlets displayed larger deviations, with measured diameters of 0.83 ± 0.07 mm (top) and 0.77 ± 0.04 mm (bottom) compared to the target value of 0.75 mm. This discrepancy may be attributed to the washing of uncrosslinked resin at the inlet interfaces, which could transiently increase local pressure and induce channel dilation. Additionally, despite employing a gelatin-based resin subjected to thermal cycling, TVAM-in-a-chip is not susceptible to the “Pandoro effect” – distorsion arising from the formation of vertical oxygen inhibitor gradient – ^25^ as the air–resin interface is eliminated through tubing clamping **(Figure 1C-ii, 4A)**.

**Figure 4.**
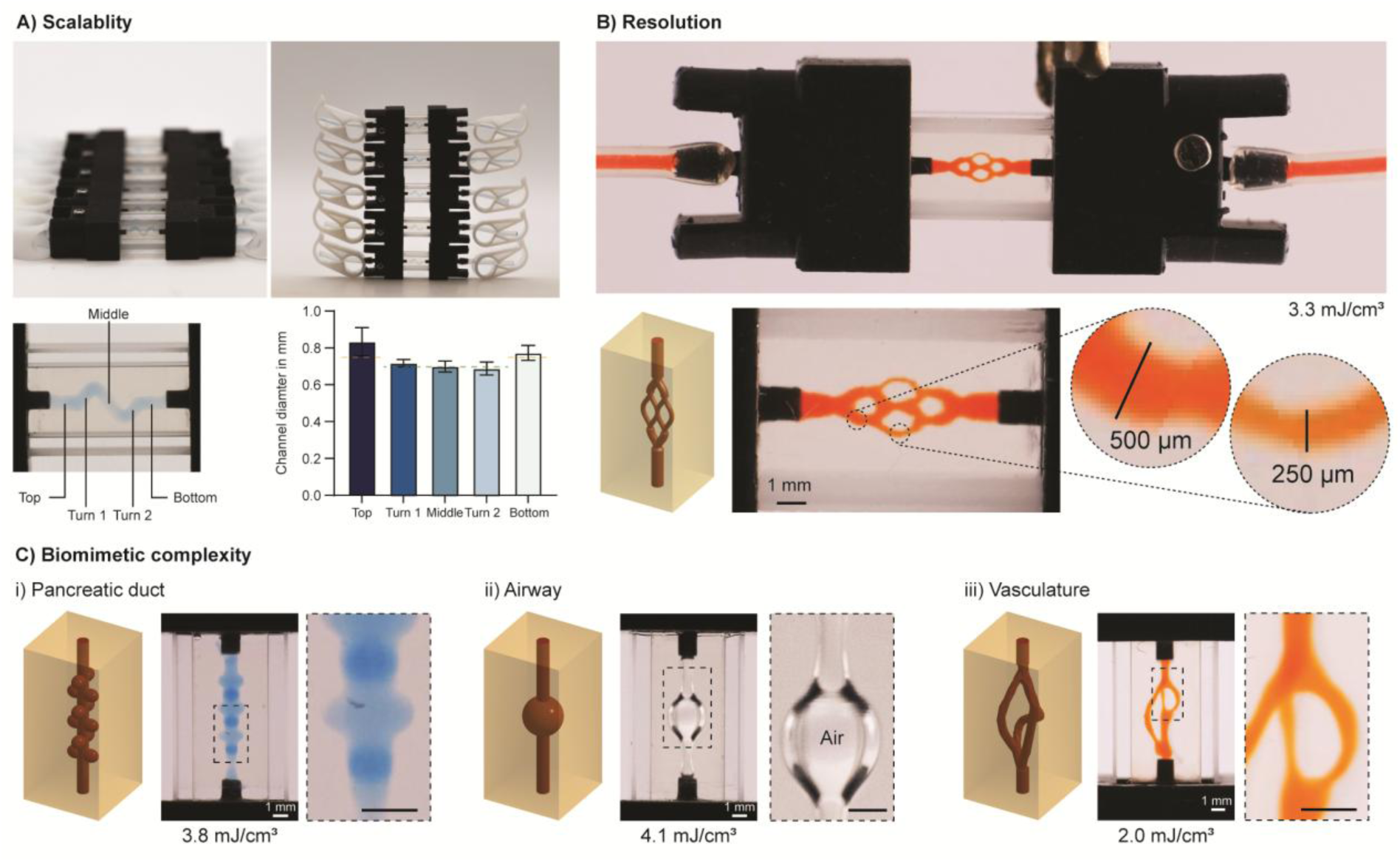
TVAM-in-a-chip resolution and complexity. (A) Representative images of five chips fabricated consecutively using the same light dose (top), along with channel diameter measurements at different positions, demonstrating the scalability and reproducibility of the approach. Dashed orange (0.75 mm) and green (0.7 mm) lines overlaid on the histogram bars indicate the respective target dimensions. (B) Branching model printed in the chip and perfused with red-dyed solution. Close-ups show the size of smaller channels down to 500 µm and 250 µm. (C) 3D models (left) and actual prints perfused with colored solutions or air of pancreatic duct (i), airway (ii), and biomimetic vasculature (iii).

Beyond reproducibility, achieving adequate resolution is essential to meet current OoC standards. Commercial organ-on-chip devices typically feature channel dimensions ranging from approximately 200 µm to 1000 µm. To benchmark the resolution of our approach, we designed a branching architecture with perfusable channels spanning 750 µm, 500 µm, and 250 µm. As displayed in **Figure 4B**, TVAM-in-a-chip can meet current standards by providing a channel resolution down to 250 µm. Notably, in soft-lithography-based platforms these channels present a non-physiological square cross section and are laying on a single plane. In contrast, TVAM-in-a-chip intrinsically provides extensive flexibility in the 3D space and provides biomimetic rounded channels. To further highlight this aspect, we generated chips mimicking pancreatic exocrine gland, consisting of a duct with acinar structures (**Figure 4C-i**), and airway system (**Figure 4C-ii**). We also implemented, in our workflow, the use of the model-driven design platform recently proposed by Sexton et al.^26^ A rapidly designed biomimetic vasculature tree .stl file was subtracted to the printable volume (5 x 5 x 10 mm), then TVAM patterns were optimized as previously described, and printed within the miniaturized chip (**Figure 4C-iii**). Importantly, these examples also showcase how a single “universal” chip platform could be exploited to generate a variety of different models.

### 2.5. 3D biomimetic platforms for perfusion culture

Of great importance for multi-day perfusion cell culture is the leak-proof integrity of the system. This remains a common limiting factor in complex organ-on-chip (OoC) platforms, particularly in 3D bioprinted models, where poor connections between inlets and the construct often arise from post-printing chip assembly **(Figure 1A)**.

In contrast, TVAM-in-a-chip enables a seamless integration between the inlet and the hydrogel, ensuring robust leak-proof performance at flow rates of up to 5 mL min^-1^, approximately 2 to 4 orders of magnitude higher than those typically used in OoC culture **(Movie S1)**.

As proof-of-concept, we aimed at generating epithelialized and endothelialized models under dynamic culture. To generate thesemodels we exploited the bioactive and high-performance 5% Gel-SH/NB photoresin. Importantly, prior to the rapid (∼ 25 s) TVAM process, the chips were sterilized (70% EtOH), preassembled and sealed in a biosafety cabinet. The pre-integrated tubing connectors facilitated efficient removal of uncrosslinked resin, subsequent channels coating and cell seeding, and reliable interfacing with the perfusion system (**Figure 5A**).

**Figure 5.**
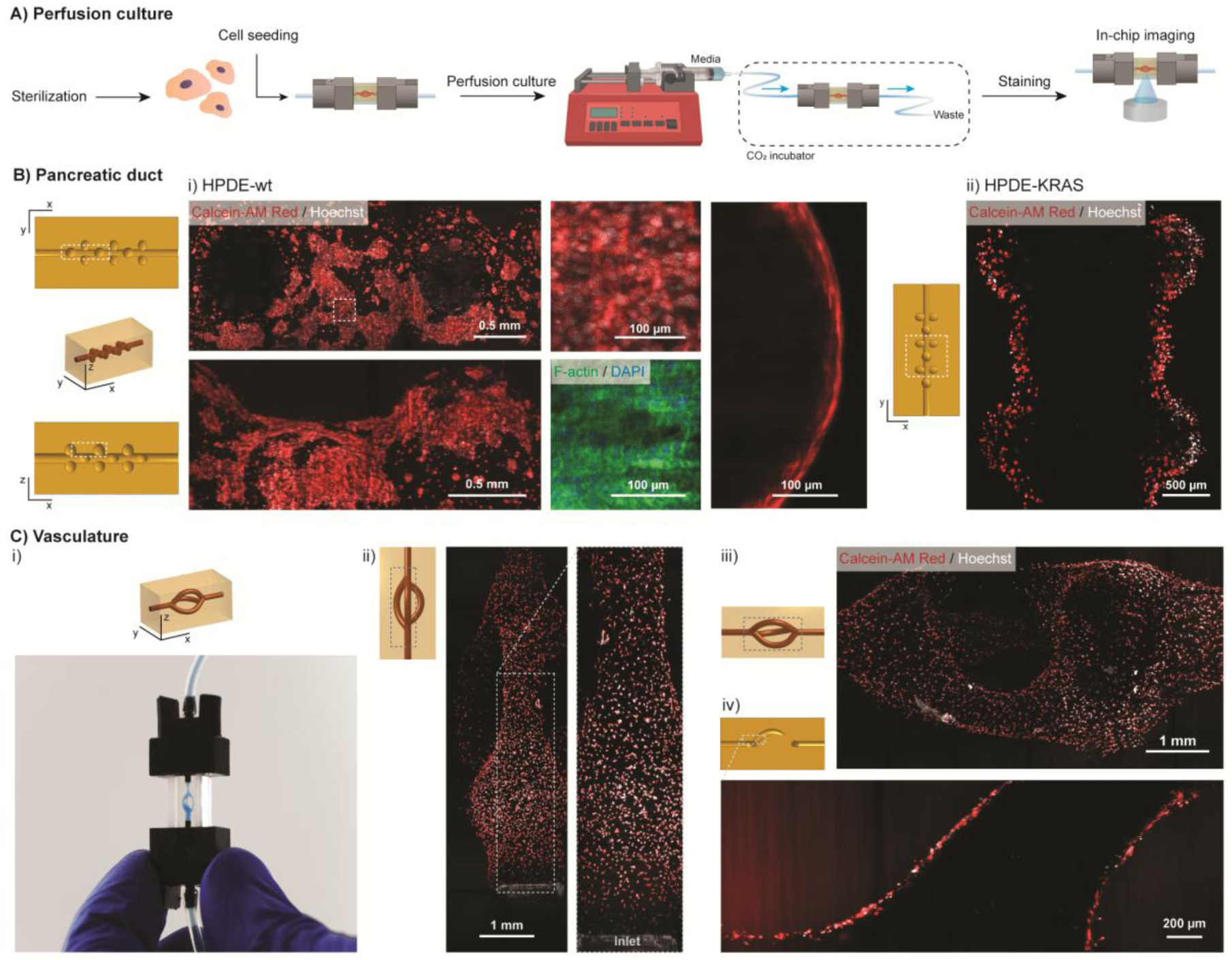
Examples of epithelialized and endothelialized TVAM-in-a-chip models. (A) Schematic of TVAM-in-a-chip steps of cell seeding, perfusion culture and imaging procedures. (B) Epithelialization of a pancreatic duct model with acinar cavities using wild type (HPDE-wt, i) and KRAS-overexpressing (HPDE-KRAS, ii) human pancreatic epithelial ductal cells. Confocal images with live (calcein-red, Hoechst) and phalloidin / DAPI stainings showing growth of monolayer epithelial patches. (C) Endothelialization of a vascular-like model (i). Confocal images with live (calcein-red, Hoechst) stainings showing successful cell adhesion (ii-iii) and localized formation of monolayers (iv).

For the epithelial example, we used wild type primary human pancreatic ductal epithelial (HPDE-wt) cells. After the seeding step (6 h), the microfluidic chip was connected to a syringe pump and cultured under a constant low flow (0.5 µL min^-1^) for 5 days. In-chip live staining and confocal imaging showed growth of live patches of epithelial monolayer covering the construct inner surface walls (**Figure 5B-i**). Additionally, F-actin staining steps were performed directly within the chip by perfusing the required solutions (e.g., fixation, permeabilization, blocking, and staining solutions) and further highlighting the presence of pancreatic epithelium growing on the inner walls of the model. Additionally, we performed a similar experiment with HPDE overexpressing the oncogene KRAS (HPDE-KRAS). HPDE-KRAS cells colonize the model showing a very distinct, rounded shape mesenchymal-leaning phenotype with limited spreading compared to the cobblestone monolayer patches reported for HPDE-wt (**Figure 5 B-ii**).

For the endothelial example, we adopted a vascular-like model (**Figure 5C-i**) seeded with human umbilical vein endothelial cells (HUVECs). After culturing the chip under constant flow (2 µL min^-1^) for 7 days, live staining showed cell adhesion across the whole construct (**Figure 5C-ii,iii**) and localized monolayer coverage (**Figure 5C-iv**).

## 3. Discussion

In this study we have presented a versatile approach, termed TVAM-in-a-chip, to attain biomimetic 3D OoCs by printing virtually any design within “universal” pre-assembled, microfluidic-ready and imaging-compatible chips. Based on rapid TVAM process and Dr.TVAM simulation software, this technology offers a new way to make single or multichannel miniaturized tissue models with a wide variety of biocompatible matrices. In particular, we demonstrated the possibility to print with synthetic or natural derived photoresins, with step-growth and chain-growth chemistry, with Type I or Type II initiators, and within a wide range of viscosities and stiffnesses. We reported on the generation of cell-laden and multicellular prints as well as compatibility with standard imaging equipment. Finally, we demonstrated high resolution (down to 250 µm channels), 3D complexity and proof of epithelialization (pancreatic duct-like model) and endothelialization (vascular-like model) in perfusion culture setups. Altogether, these results present TVAM-in-a-chip as a unique and versatile platform to attain 3D scaffold guided morphogenesis.

While this work presents a first proof-of-concept, the full potential of this technology is far from being achieved. Improving organ mimicry of these platforms requires extensive multidisciplinary work, spanning from computational and optics improvements to design of tissue tailored photoresins and optimization of cell culture conditions.

On one hand, this technology can further benefit from enhancing printing resolution to achieve more complex, organotypic microarchitectures. First, taking the vasculature model as an example, if we aim at following Murray’s law stating that the cube of a parent vessel’s radius equals the sum of the cubes of its daughter vessels’ radii, this will require finer inlets (e.g., via µSLA) to access 50-100 µm channels within the printing window. In the current setup, with inlets of ∼ 0.8 mm, this would require around 9-12 (symmetric) branching events. Entering the ≤ 50 µm regime will also require the implementation of photochemical kinetic models in the simulation framework to account for diffusion and inhibition coming from oxygen and other chemicals (e.g., TEMPO^27^ or DMPO^11^),^28^ similarly to what recently presented for single-view holographic setups.^29^

On the other hand, we envision exploring more complex biological models leveraging cell-laden photoresin formulations, which will require novel, physical-based scattering correction frameworks in Dr.TVAM. In addition, TVAM-in-a-chip appears a unique tool to elevate to 3D the promising results obtained with advanced organoid-derived 2.5D chips.^30,31^ Importantly, the promising adhesion but limited confluency of HUVECs and HPDEs—latter cells well known to adhere poorly to gelatin due to distinct integrin–ECM requirements^32–34—highlight^ that increasing biological complexity necessitates careful fine-tuning of the photoresin formulations. Future work oriented to establish predictive human (patho)physiological models will require optimization of both the biochemical and biophysical properties of the matrix. As demonstrated in this work, such tuning can be achieved across a broad palette of (photo)chemical and mechanical material parameters.

Finally, we also want to point out that, as herein done for the chip inlets, one could enhance the functions of chips by introducing functional elements (e.g., sensors, actuators) without drastically impairing the printing potential. As long as the shape, position, and optical properties are defined, one can leverage the overprinting capabilities of our open-source optical framework.^11^

## 4. Conclusion

In summary, we have introduced a versatile biomanufacturing approach with far-reaching potential, termed TVAM-in-a-chip, that overcomes some key limitations of current OoC platforms, paving the way for more complex biomimetic 3D OoC platforms. This technology offers opportunities for improvement in resolution, modeling accuracy, and biological complexity. Future advancements will rely on integrating computational, optical, and material innovations to better replicate organ-level structures and functions. Overall, TVAM-in-a-chip holds strong potential to advance the development of predictive and customizable in vitro tissue systems.

## 5. Experimental Section

### Synthesis of Gelatin-Methacryloyl (Gel-MA)

Gel-MA synthesis was adapted from previous protocol. Briefly, 25 g of gelatin type A (gel strength 300) from porcine skin was first dissolved in PBS at 10% at 45 °C. When completely dissolved, 15 mL of methacrylic anhydride was added dropwise and the reaction was left to proceed for 1.5 h. Next, the reaction mixture was diluted twofold in prewarmed mQ H_2_O, and then centrifuged at 4000 rpm for 5 minutes to separate unreacted anhydride and acrylic acid. Next, 0.5 g of sodium chloride (NaCl) was added to the supernatant, and the solution was dialyzed (MWCO: 14 kDa, Roth AG) against mQ H_2_O for 4-5 days with frequent water changes and then freeze-dried. Lyophilized G-MA was stored at -20 °C prior to use. The degree of substitution (DS) was estimated to be ∼0.25 mmol of MA per gram of gelatin with ^1^H-NMR in D_2_O using internal standard 3-(trimethylsilyl)-1-propanesulfonic acid (DSS, 2H, ∼0.65 ppm) and MA protons (∼5.45 and 5.70 ppm, **Figure S2**).

### Synthesis of Gelatin-Norbornene (Gel-NB)

Gel-NB was synthesized as previously described by Rizzo et al.^35^ In short, 20 g of gelatin type A (gel strength 300) from porcine skin was first dissolved in 0.5 M pH 9 carbonate–bicarbonate buffer at 10% at 40 °C. When completely dissolved, 0.4 g of *cis*-5-norbornene-endo-2,3-dicarboxylic anhydride (CA, Chemie Brunschwig AG) was added to the reaction mixture under vigorous stirring. Every 10 min, 0.4 g of CA was added for a total amount of 2 g. Following 20 minutes after the last addition, the solution was diluted twofold in prewarmed mQ H_2_O. Next, 2 g of NaCl was added and the solution was filter-sterilized (0.2 µm) and dialyzed (MWCO: 14 kDa, Roth AG) for 4–5 days against mQ H2O at 30 °C with frequent water changes before freeze-drying. Lyophilized G-NB was stored at -20 °C prior to use. The degree of substitution (DS) was estimated to be ∼ 0.17 mmol of NB per gram of gelatin with ^1^H NMR in D2O using internal standard 3-(trimethylsilyl)-1-propanesulfonic acid (DSS, 2H, ∼0.65 ppm) and NB protons (∼6.4–6.0 ppm, **Figure S3**).

### Synthesis of Gelatin-Thiol (Gel-SH)

Gel-SH was synthesized as previously described by Rizzo et al.^36^ In short, 10 g of type A porcine gelatin (gel strength 300) were dissolved in 500 mL of 150 mM pH 4.5 MES buffer. 0.48 g of 3,3’-dithiobis(propionohydrazide) (DTPHY, Chemie Brunschwig AG) were added to the stirring solution. Once dissolved, 1.5 g of 1-ethyl-3-(3-dimethylaminopropyl)carbodiimide (EDC, Roth AG) were added to the reaction mixture and left to proceed for 24 h. Next, 3.3 g of reducing agent tris(2-carboxyethyl)phosphine (TCEP, Chemie Brunschwig AG) were added. Reduction of disulfide bonds was left to proceed for 6 h prior to addition of 1 g of sodium chloride (NaCl) and dialysis (MWCO: 14 kDa, Roth AG) against acidified (pH 4) mQ H2O for 4-5 days with frequent water changes. The G-SH solution was then sterile filtered prior to freeze-drying. Lyophilized G-SH was stored at -20 °C prior to use. The degree of substitution (DS) was estimated to be ∼ 0.26 mmol of SH per gram of gelatin with ^1^H-NMR in D_2_O using internal standard 3-(trimethylsilyl)-1-propanesulfonic acid (DSS, 2H, ∼0.65 ppm) and hydrazide methylene protons (∼2.7 and 2.8 ppm, **Figure S4**).

### Photoresins preparation

The photoresins were prepared by dissolving the polymers in pH 7.4 PBS. 5 kDa 4-arm PEG (PEG4-MA) was purchased from Creative PEGworks, while methacryloyl hyaluronic acid (HA-MA) was purchased from Advanced BioMatrix, Inc.. For Gel-SH/NB and Gel-MA, the process was conducted at 40 °C. The photoinitiator lithium phenyl-2,4,6-trimethylbenzoylphosphinate (LAP) was used at 0.05% and added from 2.5% stock solution in PBS. The type II bimolecular photoinitiator composed of tris(2,2′-bipyridine)ruthenium(II) and co-initiator triethanolamine (Ru / TEOA) were used at 0.5 mM and 5 mM and added from 50 mM and 500 mM stock solutions in PBS, respectively. All resins were filtered (0.22 µm) prior to printing to remove scattering debris. Absorption at 399 nm was measured on a spectrophotometer (Cary 60, Agilent). Refractive index was measured with an AR200 (Reichert Inc.) digital refractometer. For gelatin-based resins, measurements were performed when in thermal gel state.

### Chip design, manufacturing and assembly

The caps were designed in Onshape (PTC). Details on the dimensions can be found in **Figure S1**. Caps were printed with SLA (Formlabs 3B, Formlabs) using the biocompatible BioMed black resin (Formlabs) and following post-processing instructor procedures. To avoid clogging of the inlets, each channel was injected to with isopropyl alcohol to remove excess of uncrosslinked resin prior to post-curing. The 5 mm quartz cuvettes (QC2001, CostLab) were cut into 20 mm sections using a diamond wire saw. The chips were assembled by simply press-fitting the caps onto the cuvette sections. For each chip, only one of the two caps featured a venting hole to evacuate air bubbles. Once closed, the chip was connected to microfluidic silicone tubing (Saint-Gobain Silicone Tubing, Diameter Inner: 0.8 mm, Diameter Outer: 4 mm, Thickness: 0.8 mm) using the barbed adapters. The photoresin (at 37°C for gelatin based resins, and room temperature for others) was then injected in the chip via the tubing using a 1 mL syringe. As shown in **Figure 1C-ii**, each side was first primed with resin to avoid bubble formation, and the filling was then performed on the side featuring the venting hole. Lastly, the venting hole was sealed using a 3 mm M1.6 screw.

### (Photo)rheology

Photorheology was carried out on an Anton Paar MCR102 rheometer equipped with an 25 mm stainless steel parallel plate and quartz glass floor. Omnicure Series1000 lamp (Lumen Dynamics) with 400 nm filter (FB400-40, Thorlabs) was used as light source. Light intensity was tuned to be 10 mW cm^-2^ at the sample location. Intensity was measured using a photodiode power sensor (S140C, Thorlabs). The photoresins were prepared as previously stated. Oscillatory measurements were performed in triplicate using 110 μL of photoresin with a 200 μm gap, at 1 Hz frequency, and 1% strain. All tests were conducted in the dark, and in the presence of a water-soaked tissue in the chamber to prevent samples from drying. Tests with gelatin-based resins were conducted at 37 °C to avoid thermal gelation. All other tests were conducted at 22 °C. Light irradiation was started after 120 s. Viscosity was measured using the same setup with flow sweep tests. Shear rate ranging from 1 to 10 s^-1^ was investigated. Reported values refer to 1 s^-1^.

### TVAM setup

TVAM is performed using a custom built setup, as previously described (see also **Figure S5**).^19^ In short, as light source we used a High Power Laser 399 nm 8 W Fiber Coupled Laser (CivilLaser, China). The high-speed digital micromirror device (DMD) (VIS-7001, Vialux), then projects the laser beam to the priting plane via a 4f system (150 mm and 100 mm plano-convex lens) with a Fourier stop to filter out higher diffraction orders. The pixel size at the image plane was measured to be 20.3 x 20.3 µm. The DMD is synchronized with the high precision rotation stage (X-RSW60C, Zaber) via an Arduino Nano Every.

### Pattern optimization and light dose calculation

All 3D models were designed in Onshape (PTC). The biomimetic vasculature of **Figure 4C-iii** was generated using the recently published algorithm by Sexton et al.,^26^ and subtracted to a cuboid of 5 x 5 x 10 mm (chip size) using Onshape. Pattern optimization was conducted as previously described^18,19^ on a NVIDIA L40S GPU. Optical properties of the chip components (cuvette and resins) used in the configuration files can be found in **Table S1-2**. All optimizations were performed with 600 patterns, a lower threshold (t_l_) of 0.6, an upper threshold (t_u_) of 0.9, a weight sparsity term of 0.02 and 40 optimization steps (**Figure S6**). An alternative pattern optimization not accounting for overprinting scenario and multi-step printing process was also adopted when discrepancies between target and chip size occurred (**Figure S7**). For example, a slightly longer cuvette section would result in misalignment between projection and chip edges and inlets, and thus print failure. When for each print the optimal printing time and laser power were identified, the target volumetric light dose in mJ cm^-3^ was calculated as follows. The projection system delivers fixed optical power. The DMD scales the projected pattern such that the brightest pixel is driven at full power, and all other pixels are normalized relative to this maximum. This normalization allows us to determine the total optical power actually delivered to the build volume. Because TVAM intentionally prescribes a three-dimensional dose distribution, knowledge of the incident power enables us to quantify how the input power is partitioned among (i) absorption within the target volume, (ii) absorption in non-target (void) voxels, and (iii) transmitted power. The absorbed power in the target can then be integrated over the exposure (printing) time to yield the absorbed energy dose (in case of constant power simply multiplied by printing time). Finally, dividing this absorbed energy by the absolute target volume provides an energy-per-volume dose metric.

### Printing process

The chip, assembled as described above, was mounted onto the rotation stage. Its angular orientation was then aligned by matching reflection of a reference red laser positioned orthogonal to the printing axis, previously calibrated from back-reflected signal from printing laser. A camera positioned along the printing axis, behind the chip and a ND filter, was instead used to register the chip position along the z axis. A vertical shift of the pattern within the DMD was used to match the chip position in z. The pre-calculated patterns were then projected to the rotating vial and the printing process was stopped upon evident change in refractive index (monitoring via camera). All prints were conducted with a max laser intensity (white image projection) between 25 and 100 mW cm^-2^. The printing process was repeated between 3-5 times to identify the optimal dose. For gelatin-based formulations, the photoresin was left to thermally gel for 5-15 minutes prior printing. After printing, the same chips were placed in a 37°C oven for a few minutes to melt the uncrosslinked resin which was then washed out with pre-warmed PBS injected via the tubing.

### Cell culture

Human foreskin fibroblasts (HFF-1 SCRC-1041, ATCC), wild type human pancreatic ductal cells (HPDE-wt) and KRAS overexpressing HPDE (HPDE-KRAS) were cultured in Dulbecco’s Modified Eagle’s Medium (DMEM) without phenol red, supplemented with antibiotic-antimycotic (Gibco), 2% L-glutamine (Gibco) and 10% fetal bovine serum (Gibco). Media was changed every other day, and cells were used at passage 6-12. HPDE cells were kindly provided by Prof. Bussolino (Candiolo Cancer Institute, FPO—IRCCS, Candiolo, Italy). Human umbilical vein endothelial cells (HUVECs, Lonza) were cultured in EGM-2 (Lonza) with media change every other day. Cells were expanded and used for the reported experiments at passage 2-5.

### Leak test

Spiral chip was generated as previously reported using 5% Gel-SH/NB photoresin. After gentle washing of uncrosslinked resin from the channel, the chip was connected to a syringe pump (PHD ULTRA, Harvard apparatus). A solution of PBS with red dye or blue-dextran was then pumped at flow rates ranging from 0.1 mL min^-1^ to 5 mL min^-1^ **(see Movie S1)**.

### Chips epithelialization and endothelialization

Chip components (tubes, caps, cuvettes, screws) were sterilized in 70% EtOH for 30 min, left to dry in a biosafety cabinet, and then assembled. 5% Gel-SH/NB photoresin was prepared as previously stated, sterile filtered (0.22 μm Millex®-GP, Millipore Sigma) and used to fill the preassembled sterile chips. Tubes were then clamped and the resin left to gel at room temperature for 10-15 minutes. TVAM process was performed as indicated above, after which the sterile chips were transferred to a humified CO_2_ incubator at 37 °C. After 10-15 minutes, warm cell culture media was carefully injected on one side to remove the uncrosslinked resin using a 1 mL syringe. The chip was left to equilibrate overnight in a humified CO_2_ incubator at 37 °C. A collagen solution (125-50, Sigma Aldrich) was then injected, and the coating process was left to occur in the incubator for 30 min. A 15 million cells mL^-1^ (HPDE and HPDE-KRAS) or 6 million cells mL^-1^ (HUVEC) suspension in the respective media was then injected into the constructs. Seeded chips were then placed in the incubator and turned every 5 minutes (0°, 180°, 90°, 180°), followed by the same process every 20 minutes to ensure homogeneous cell adhesion. Cells were left to adhere for 6 h prior to connecting the chips to a syringe pump (PHD ULTRA, Harvard apparatus). 20 mL (HPDE) or 50 mL (HUVECs) syringes filled with media were used for the perfusion culture until the experiment was stopped (5 days at 0.5 µL min^-1^ for the pancreatic model, and 7 days at 2 µL min^-1^ for the vasculature model). The pump was placed outside the incubator. Around 50 cm of tubing inside the incubator ensured temperature and gas equilibration of the media prior to entering the chip. The outlets of the chips were connected to a waste container.

### Imaging

Confocal imaging was performed on an inverted confocal microscope (Leica SP8) equipped with a HC PL Fluotar 10x/0.30 objective (11 mm WD). Pictures of the chip and printed constructs were obtained with a Fujifilm XT-3 camera.

## Acknowledgements

The authors thank Claude Amendola and the EPFL mechanical workshop for the cutting of the quartz cuvette. The authors also thank Prof. Chiara Tonda-Turo from Politecnico di Torino for providing HPDE-wt and HPDE-KRAS cells.

## Data Availability Statement

All data are available in the main text, the supplementary materials. Additional data can be found at: https://github.com/EPFL-LAPD/Tomographic-Printing-in-a-Chip-A-Versatile-Platform-for-Biomimetic-3D-Organ-on-Chip

