## Supplementary Information for "Tomographic Printing in a Chip: A Versatile Platform for Biomimetic 3D Organ-on-Chip"


**This PDF file includes:**

Figure S1 to S7
Movie S1
Tables S1 to S2

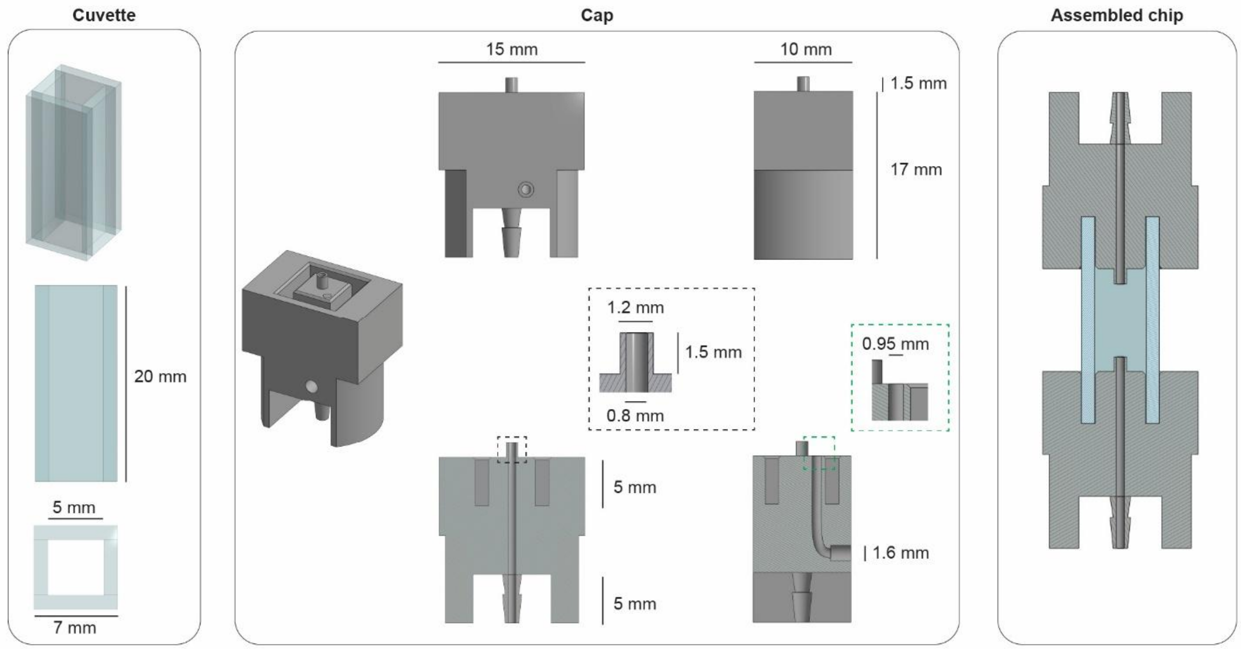


**Figure S1.** Details of chip design.

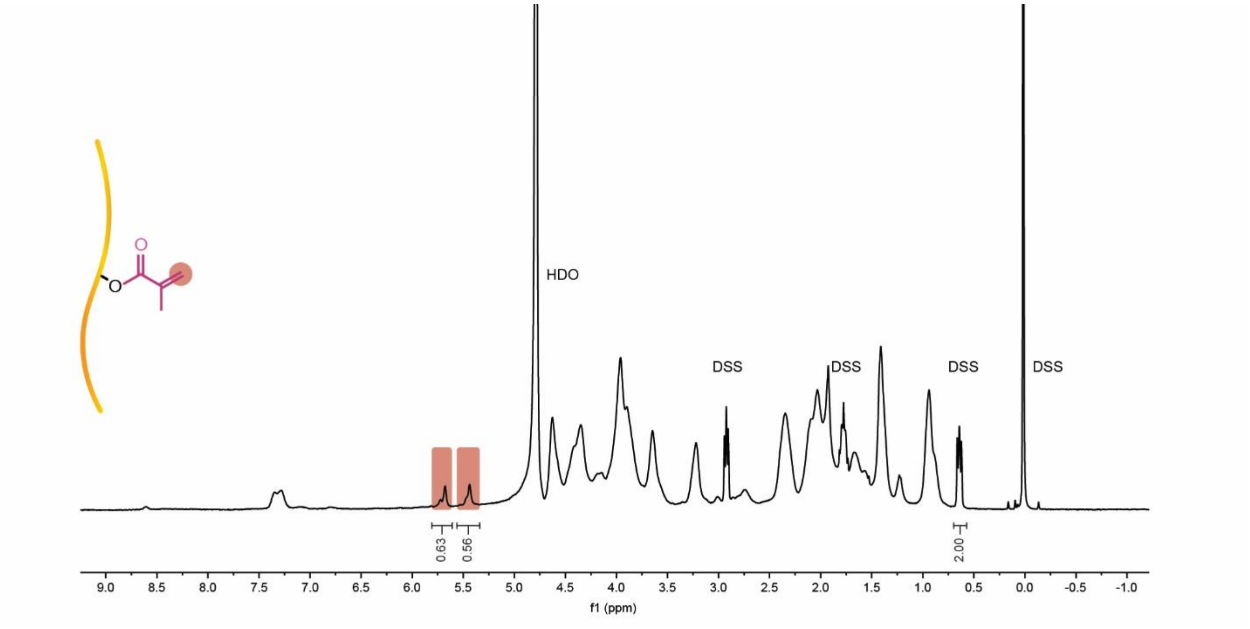


**Figure S2.** Gel-MA $^{1}$H-NMR.

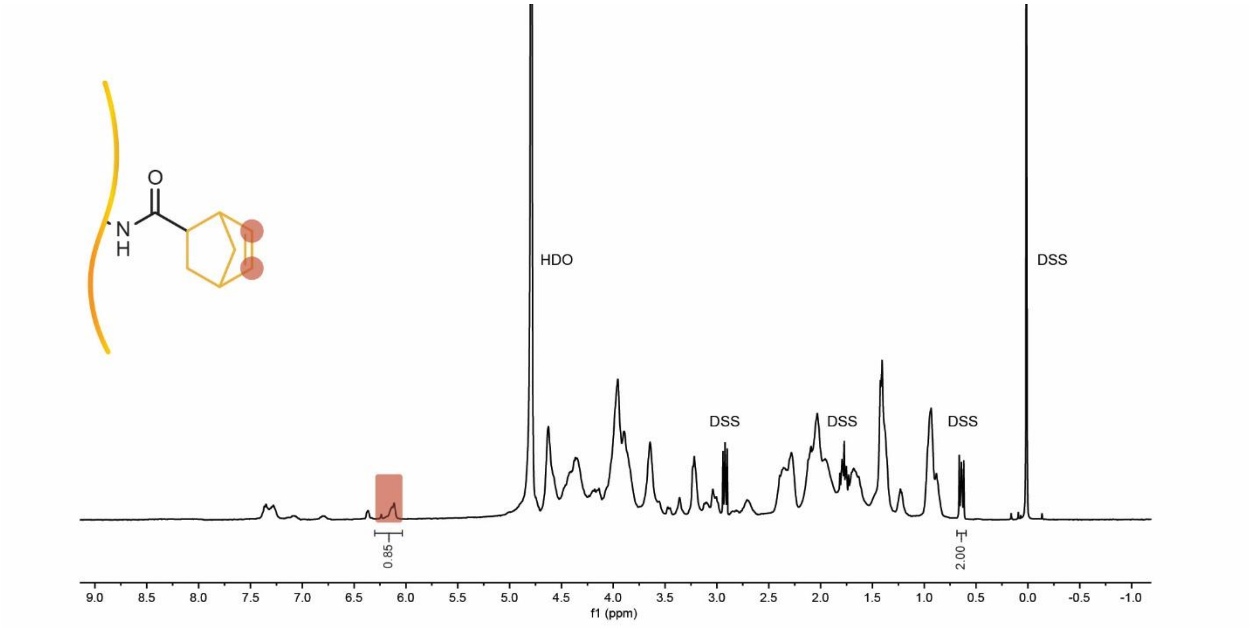


**Figure S3.** Gel-NB $^1$H-NMR.

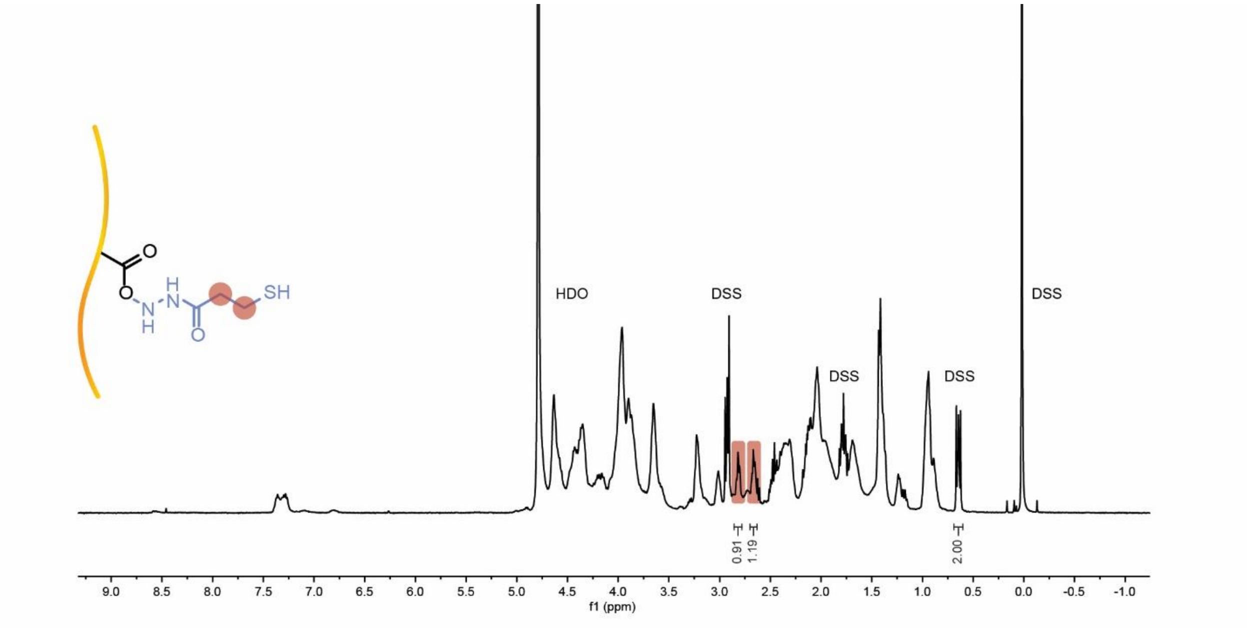


**Figure S4.** Gel-SH $^{1}$H-NMR.

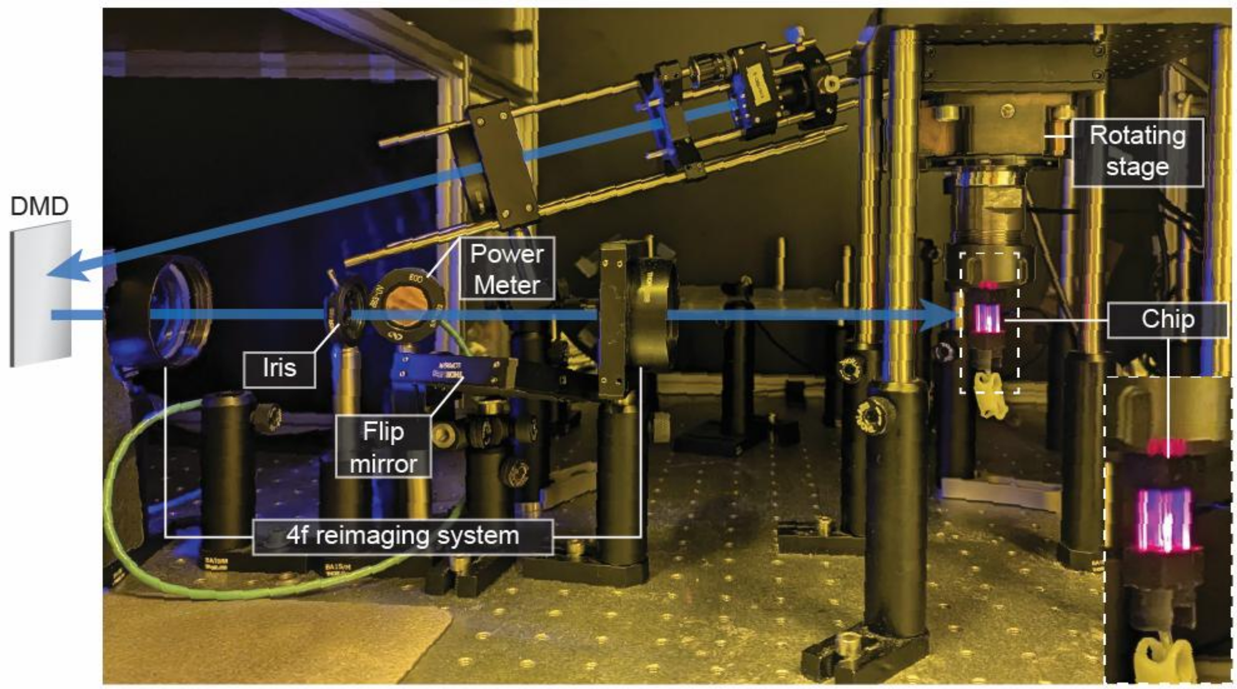


**Figure S5.** TVAM setup.

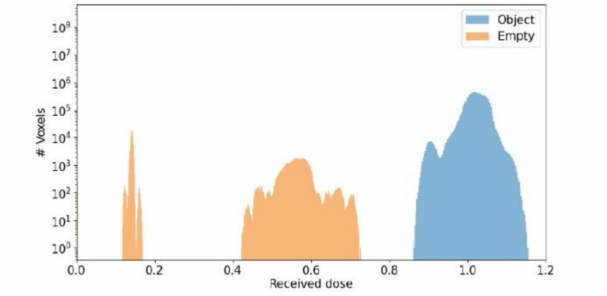


**Figure S6.** Example of Dr.TVAM output histogram reporting intensity values of the empty (uncrosslinked) regions in orange, and intensity values of the object (crosslinked) regions in blue. The good separation between the two is a sign of successful optimization and consequently likely successful print. Note that the good separation also comes from the upper ($t_u$) and lower ($t_l$) threshold defined in the configuration file (see Table S2), forcing dose in empty regions to stay below $t_l$ and dose in object regions to stay above $t_u$. For more information visit the Dr.TVAM documentation: https://drtvam.readthedocs.io/en/latest/basic_usage.html .

.

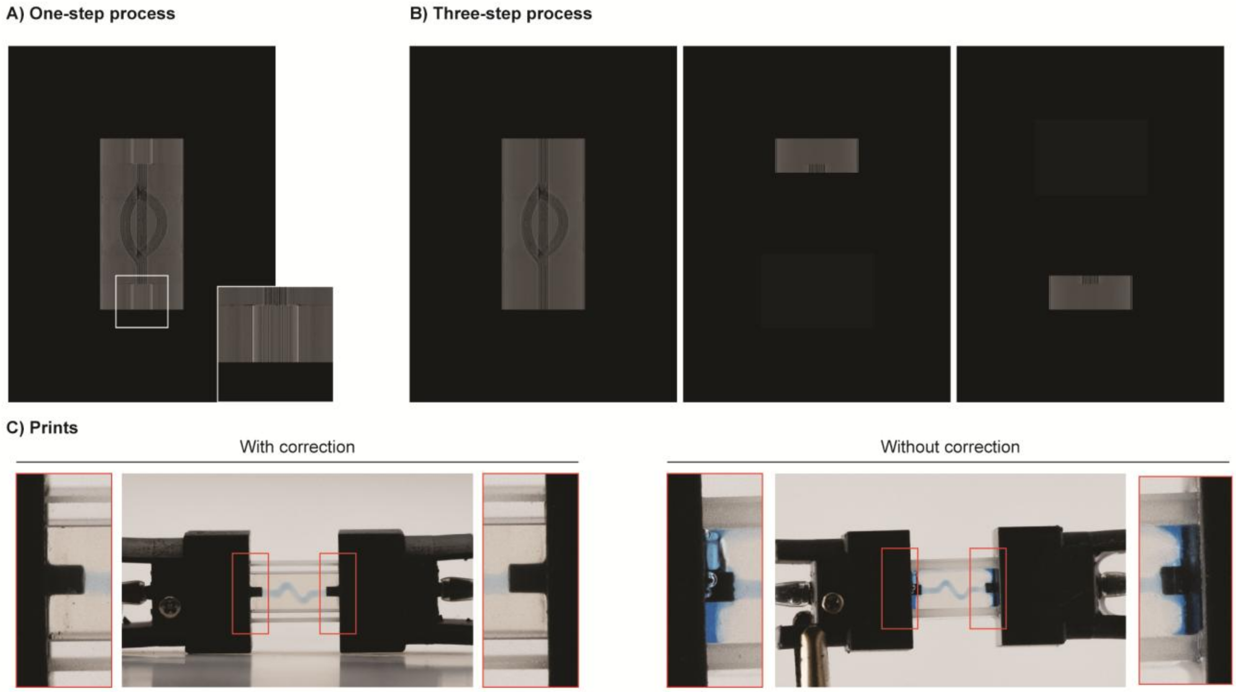


**Figure S7.** A) Example of pattern at 0º with overprinting features. The optimization accounts for the presence of light occluding elements (inlet / outlet) and delivers more light in those regions (close-up). This approach enables a T-OoC in a single printing process. B) Example of pattern at 0º not accounting for light occluding elements (left). As a result of such print, the top and bottom parts of the chip remain underpolymerized and require further illumination (middle, right) in a multi-step process. While more time-consuming, this approach is effective when the cuvette cross-section or cap size deviates from the values used for overprinting optimization, preventing alignment issues. C) Examples of print with (left) and without (right) overprinting strategies showing underpolymerization and leakage when not accounting for light occluding inlets.

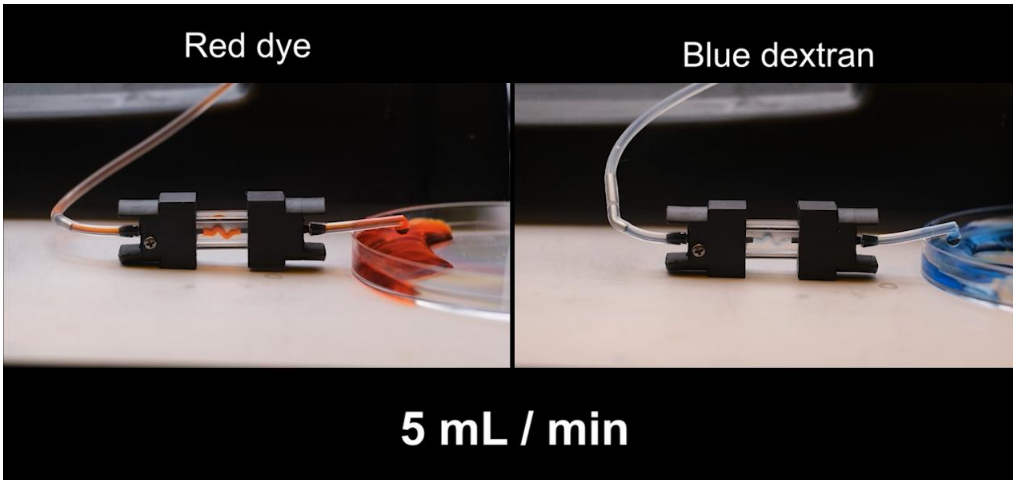


**Movie S1.** Leak test for TVAM-in-a-chip system. No leakage is observed across flow rates ranging from 0.1 mL $min^{-1}$ to 5 mL $min^{-1}$, approximately 2 to 4 orders of magnitude higher than those typically employed in OoC culture.

**Table S1.** Photoresin optical properties and printing parameters.

| Formulation | Refractive Index | Attenuation coefficient | Target | Max intensity* in mW $cm^{-2}$ | Printing time in s | Target absorbed dose in mJ $cm^{-3}$ |
|---|---|---|---|---|---|---|
| PEG4-MA 20%, LAP 0.05% | 1.358 | 0.1103 | Spiral | 100 | 98.4 | 31.1 |
| HA-MA 2%, LAP 0.05% | 1.3374 | 0.0401 | Spiral | 100 | 41.6 | 11.6 |
| Gel-MA 10%, LAP 0.05% | 1.3512 | 0.0609 | Spiral | 50 | 42.4 | 6.3 |
| Gel-MA 10%, 0.5 mM Ru 5 mM SPS | 1.3506 | 0.2216 | Spiral | 100 | 41.6 | 15.8 |
| Gel-SH/NB 5%, LAP 0.05% | 1.345 | 0.0609 | Spiral | 50 | 25.6 | 3.8 |
| Gel-SH/NB 10%, LAP 0.05% | 1.3517 | 0.0609 | Spiral | 50 | 14.4 | 2.1 |
| Gel-SH/NB 15%, LAP 0.05% | 1.3617 | 0.0609 | Spiral | 50 | 10.4 | 1.5 |
| Gel-SH/NB 5%, LAP 0.05% | 1.3517 | 0.0609 | 1-channel | 50 | 23.9 | 3.6 |
| Gel-SH/NB 5%, LAP 0.05% | 1.3517 | 0.0609 | 2-channels | 50 | 23.4 | 3.4 |
| Gel-SH/NB 5%, LAP 0.05% | 1.3517 | 0.0609 | 3-channels | 50 | 25.0 | 3.7 |
| Gel-SH/NB 5%, LAP 0.05% | 1.3517 | 0.0609 | 2-0 channel | 50 | 24.96 | 3.6 |
| Gel-SH/NB 5%, LAP 0.05% | 1.3517 | 0.0609 | 2-1 channels | 50 | 28.1 | 4.2 |
| Gel-SH/NB 5%, LAP 0.05% | 1.3517 | 0.0609 | 3-1 channels | 50 | 24.96 | 3.6 |
| Gel-SH/NB 5%, LAP 0.05% | 1.345 | 0.0609 | 1-channel cell-laden | 50 | 23.9 | 3.6 |
| Gel-SH/NB 5%, LAP 0.05% | 1.345 | 0.0609 | Branching [Fig. 3A] | 50 | 22.4 | 3.3 |
| Gel-SH/NB 5%, LAP 0.05% | 1.345 | 0.0609 | Pancreatic model | 50 | 25.6 | 3.8 |
| Gel-SH/NB 5%, LAP 0.05% | 1.345 | 0.0609 | Airway model | 50 | 27.2 | 4.1 |
| Gel-SH/NB 10%, LAP 0.05% | 1.3517 | 0.0609 | Vasculature model [Fig. 4] | 25 | 27.2 | 2.0 |
| Gel-SH/NB 5%, LAP 0.05% | 1.345 | 0.0609 | Vascular-like model [Fig. 5] | 50 | 25.6 | 3.8 |
| All prints are performed at a rotational speed of 4 s per turn, and 600 patterns per rotation. *Max intensity: light intensity when all DMD pixels are turned on (255 greyscale values). | | | | | | |

**Table S2.** Example of Dr.TVAM JSON configuration file for pattern optimization. For more information visit the Dr.TVAM documentation:
https://drtvam.readthedocs.io/en/latest/basic_usage.html .

| | |
|---|---|
| { | |
| "vial": { | // defines container and photoresin |
| "type": "custom", | |
| "filename_vial_outer": "cuvette_outer.ply", | // ply files of cuvette outer shell |
| "filename_vial_inner": "cuvette_inner.ply", | // ply files of cuvette inner shell |
| "ior": 1.4702, | // refractive index of glass cuvette |
| "medium": { | // photoresin |
| "ior": 1.3517, | // refractive index of photoresin |
| "phase": { | |
| "type": "rayleigh" | |
| }, | |
| "extinction": 0.0609, | // extinction coefficient of photoresin |
| "albedo": 0.0 | // albedo is 0, so no scattering |
| } | |
| }, | |
| "filter_corner": { | //exclusion of cuvette corners from optimization |
| "dist": 3.735, | |
| "radius": 1.277 | |
| }, | |
| "projector": { | // DMD |
| "type": "collimated", | // type of light source (collimated laser) |
| "n_patterns": 600, | // number of desired projection patterns |
| "resx": 768, | // number of x pixels |
| "resy": 1024, | // number of y pixels |
| "cropx": 650, | // number of x pixels used in the optimization |
| "cropy": 650, | // number of y pixels used in the optimization |
| "crop_offset_y": 187, | // offset the crop area in y to center the patterns |
| "crop_offset_x": 59, | // offsets the crop area in x to center the patterns |
| "pixel_size": 20.3e-3, | // pixel size at the printing plane |
| "motion": "circular", | // motion of the vial |
| "distance": 20 | // distance of emitter to vial (irrelevant for collimated) |
| }, | |
| "sensor": { | // 3D grid where absorption measurements are taken |
| "type": "dda", | |
| "scalex": 13.195, | // physical size of absorption region |
| "scaley": 13.195, | // physical size of absorption region |
| "scalez": 13.195, | |
| "film": { | // sensor discretization in space |
| "type": "vfilm", | |
| "resx": 650, | // number of voxels in each dimension |
| "resy": 650, | |
| "resz": 650 | |
| } | |
| }, | |
| "target": { | // object to be printed |
| "filename": "target.ply", | |
| "size": 10 | // size in mm |
| }, | |
| "loss": { | // loss function parameters |
| "type": "threshold", | |

| | |
|---|---|
| "tl": 0.6, | // lower threshold |
| "tu": 0.9, | // upper threshold |
| "weight_sparsity": 0.02, | // weight of sparsity term. The higher the value, the lower the sparsity resulting in higher pattern efficiency and lower contrast. |
| "M": 4 | // power to which to raise the pattern values |
| }, | |
| "filter_radon": false, | |
| "spp_ref": 24, | // light paths per proj. pixel when evaluating final results |
| "spp": 24, | // light paths per proj. pixel in forward model |
| "spp_grad": 24, | // light paths per proj. pixel in backpropagation model |
| "transmission_only": false, | // transmission and reflection |
| "n_steps": 40, | // optimization steps |
| "output": "/location path" | |
| } | |
